# Dispersal syndrome evolution in angiosperms: ancestral zoochory, correlations with pollination mode and biome, and sampling effects

**DOI:** 10.1101/2025.05.26.656093

**Authors:** Dieder de Frens, Freek T. Bakker, Hervé Sauquet, Ruby E. Stephens, Renske E. Onstein

## Abstract

**Aim:** Angiosperms have evolved a diverse range of seed dispersal syndromes that have undoubtedly played a key role in their evolutionary history and rise to dominance. However, it remains largely unknown how ancestral angiosperm seeds were dispersed, whether dispersal syndromes are associated with biome and pollination mode transitions, and how sampling and syndrome assignment affect historical inferences at the angiosperm-wide scale.

**Location:** Global

**Time period:** Cretaceous - Cenozoic

**Major taxa studied:** Angiosperms

**Methods:** We integrated a large phylogeny with a high quality dispersal-related trait and syndrome dataset for >1200 species representing all extant angiosperm families. We used phylogenetic comparative methods to infer ancestral states of major angiosperm clades and evolutionary correlations between dispersal syndromes, biomes and pollination modes. In addition, we used taxon removal simulations to assess how sampling affects ancestral state and transition rate inferences.

**Results:** We show that angiosperms were ancestrally most likely animal-dispersed (zoochory), with fleshy, indehiscent fruits. Over macroevolutionary time, frequent transitions between dispersal syndromes occurred, with non-specialised (autochorous) dispersal as an intermediate state between transition from animal (zoochory) to wind (anemochory) dispersal. Furthermore, we found that zoochory was associated with increased evolutionary transitions to tropical biomes and to biotic pollination, and abiotic dispersal to cold or arid biomes and to abiotic pollination. Taxon removal up to 50% showed minimal effects on ancestral state and transition rate inferences.

**Main conclusions:** The progressive evolution of diaspore complexity may have facilitated the frequent and repeated transition between dispersal syndromes, critical for the colonisation and exchange of angiosperms across modern biomes, and present-day associations between biotic and abiotic pollination and seed dispersal.

## Introduction

Since their first emergence between 140 and 270 million years ago (Mya) (Sauquet et al., 2022), angiosperms have quickly become evolutionarily successful both in terms of species richness – with ca. 370,000 species alive today – and ecological dominance on land, especially compared to other major plant clades (e.g., bryophytes, ferns, and gymnosperms) (Crepet & Niklas, 2009). This rise to dominance is usually referred to as the ‘Angiosperm Terrestrial Revolution’ (ATR, from 100 to 50 Mya, Benton et al., 2022), during which most extant angiosperm families emerged and spread across the major terrestrial biomes (Li et al., 2019; Ramírez-Barahona et al., 2020, Ding et al. 2025). Many characters probably facilitated the explosive angiosperm radiation and their wide distribution, including a higher rate of character evolution (‘trait flexibility’ *sensu* Onstein, 2020), fast generation times (Smith & Donoghue, 2008), photosynthetic capacity (McElwain et al., 2016), a reduction in genome size (Simonin & Roddy, 2018), and the development of flowers, double fertilization, and fruits (Endress, 2011, Benton et al., 2022). Although seed dispersal is evidently an important process in the ecology and evolution of plants (Howe & Smallwood, 1982; Nathan et al., 2008; Beckman & Sullivan, 2023; Jin et al., 2026a), it remains unclear how dispersal mechanisms have shaped the spread of angiosperms across the major biomes or changes in their reproductive (e.g., pollination) biology.

Angiosperm fruits possess a range of functional characters that play key roles in seed dispersal. This includes fruit characters such as dehiscence (i.e., ripe fruits split to release or present the seeds) (Balanzà et al., 2016), fleshiness (i.e., whether fruits have a fleshy pericarp) (Valenta & Nevo, 2020), the presence of different types of appendages, such as wings, on either seeds or fruits, the presence or absence of air-retaining structures (Howe & Smallwood, 1982), fruit or seed colour (Nevo et al., 2018), odour (Nevo et al., 2016), and nutritional compounds (Konečná et al., 2018). Different associations of such traits constitute functional groupings known as dispersal syndromes – including abiotic (e.g., wind or water dispersal) and biotic (e.g., animal dispersal) mechanisms (Rojas et al., 2022). Interestingly, the possession of these syndromes alone is not sufficient to set angiosperms apart from gymnosperms, as most main syndromes present in angiosperms are also found in gymnosperms, with the exception of autochory (e.g., explosive dispersal) (Herrera, 1989; van der Pijl, 1982). However, the increased number of possible dispersal-related adaptations, and thus complexity and variation of their disseminules (diaspores), is unique for angiosperms. Where gymnosperms can essentially only differentiate their cones, seeds, and in some cases arils (Contreras et al., 2017), angiosperms can differentiate their pericarp layers, seeds, arils, persistent floral organs, and bracts, thereby possibly increasing their adaptability to different selection pressures. Because of this variability, dispersal syndromes can convergently evolve by modifying different features or ‘traits’ (**Fig. 1**).

**Figure 1.**
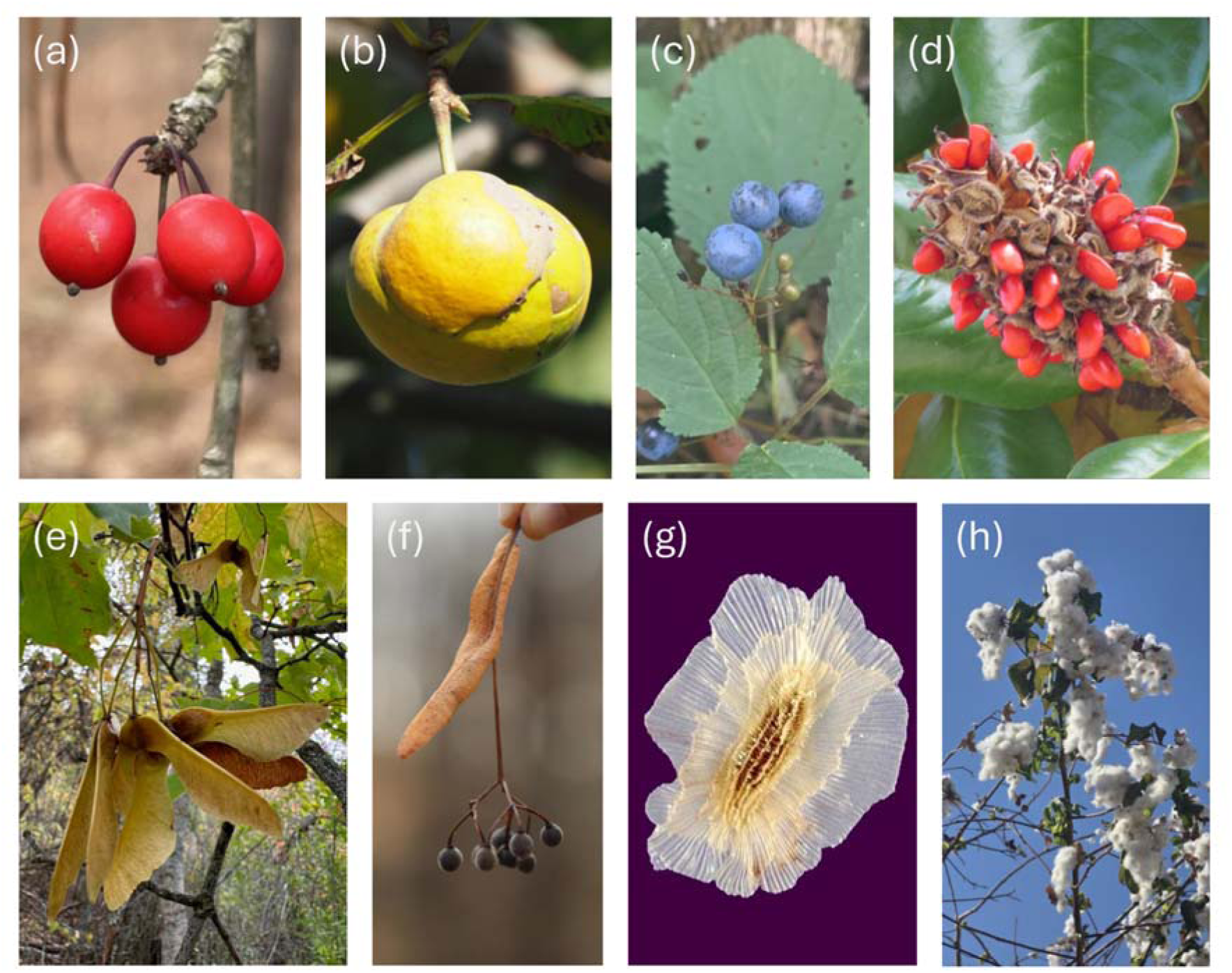
Examples of different diaspore types associated with the same dispersal syndromes in zoochorous and anemochorous species. Zoochorous examples are shown in (a–d) and anemochorous examples in (e–h), illustrating dispersal by whole fruits, floral fruits, seeds, and covered seeds, respectively (definitions in **Table S1**). Whole-fruit dispersal is represented by (a) *Ilex decidua* and (e) *Acer platanoides*; floral-fruit dispersal by (b) *Dillenia indica* and (f) *Tilia americana*; seed dispersal by (c) *Caulophyllum thalictroides* and (g) *Paulownia tomentosa*; and covered-seed dispersal by (d) *Magnolia grandiflora* and (h) *Gossypium hirsutum*.

Dispersal syndromes are not randomly distributed across biomes, time periods, and ecological settings. For example, in modern angiosperms, the proportion of fleshy-fruited and animal-dispersed (zoochorous) species is positively correlated with warmer, wetter, and lower latitude habitats (Chen et al., 2017; Correa et al., 2015; Wang et al., 2022). Fossil and phylogenetic evidence also illustrates how the frequency of dispersal syndromes has changed throughout geological time, possibly in relation to changing global environments (Eriksson et al., 2000; Vasconcelos et al., 2023; Jin et al., 2026a). However, the dispersal mechanism of early angiosperms remains ambiguous; hypothetically, small diaspore sizes and lack of biotic dispersal vectors in early angiosperm floras may be indicative of abiotic seed dispersal, possibly up until the Late Cretaceous (100.5–66 Ma) (Tiffney, 1984; Tiffney, 2004; Wing & Tiffney, 1987; Eriksson et al., 2000). Additionally, it remains unclear to what extent dispersal syndrome evolution influenced the spread of angiosperms across modern biomes (Donoghue & Edwards, 2014; Vasconcelos et al., 2023), given that dispersal ability and certain dispersal syndromes (e.g., zoochory) may facilitate long distance dispersal and range expansion (Alzate & Onstein, 2022). Similarly, pollination modes – such as by insects, vertebrates or wind – show strong associations with habitat type (open/closed) and latitude, suggesting an evolutionary association with biome and – possibly indirectly – with seed dispersal mode (Rech et al., 2016; Kriebel et al., 2019; Stephens et al., 2023; Yamawo & Ohno, 2024). For example, ca. 82% of endozoochorous trees are biotically pollinated, whereas > 80% of anemochorous trees are wind-pollinated, suggesting joint evolutionary associations between pollination and seed dispersal mutualisms (Yamawo & Ohno, 2024).

Despite the overwhelming interest in dispersal biology of angiosperms (Beckman & Sullivan, 2023), it remains unclear how ancestral angiosperm seeds were dispersed and whether dispersal syndrome evolution has been associated with biome and pollination mode transifffffons. Recent work across >35k angiosperm species showed a lack of association between diversification rates and dispersal syndromes, but a strong association between dispersal syndrome transitions and paleoclimate, and between the geographical distribution of dispersal syndromes and contemporary climate (e.g., temperature) (Jin et al., 2026a). However, their sampling did not include all angiosperm families (ca. 121 out of 418 families missing) and Amborellaceae, representing the extant sister lineage to all other extant angiosperms, was missing. This hindered a comprehensive reconstruction of ancestral dispersal syndromes across families and orders, and ambiguity in the angiosperm ancestral dispersal syndrome (Jin et al., 2026a). Furthermore, dispersal syndromes were classified based on available information from various heterogeneous sources using different labels and criteria to place plant species into seed dispersal categories, which raised concerns about data quality (Wasowicz & Green, 2026, but see Jin et al., 2026b).

Here, we address these limitations by assembling a high quality dispersal trait database and applying a trait-based quantitative framework to objectively assign species to dispersal syndromes, focusing on >1200 species that represent all extant angiosperm families. We integrated our database with a robust phylogenetic tree (Ramírez-Barahona et al., 2020), pollination mode (Stephens et al., 2023) and super biome data (Ramírez-Barahona et al., 2020) to (i) quantify ancestral dispersal syndromes and transition pathways in the evolution of dispersal-related characters and dispersal syndromes across angiosperms; (ii) assess evolutionary correlations between dispersal syndrome, pollination mode and biome evolution and (iii) evaluate data quality and sampling and their effects on ancestral dispersal syndrome and transition rate inferences. We hypothesize that (H1) the evolution of dispersal syndromes is labile, with frequent transitions and high transition rates between dispersal syndromes – i.e., animal (zoochory), wind (anemochory), water (hydrochory) and autochory (autochory, barochory, and antitelochory) – due to the multitude of characters that can convergently evolve into similar syndromes; and (H2) that dispersal syndrome evolution is correlated with shifts in angiosperm ecology, with transitions across biomes (temperate/tropical/arid) and pollination modes (biotic/abiotic) contingent on dispersal syndrome.

## Materials and Methods

### Dispersal traits and syndrome assignment

We collected dispersal-related characters for the 1201 species used in Stephens et al. (2023), which is based on the angiosperm phylogenetic tree constructed by Ramírez-Barahona et al. (2020). This tree was designed to broadly represent the diversity across all angiosperm families, with additional species from larger families. Hence, the sampled species are phylogenetically representative of the c. 370 000 species of angiosperms, but random with respect to their seed dispersal mode. In addition, biome data (Ramírez-Barahona et al., 2020) and pollination mode data (Stephens et al., 2023) were available for the exact same set of species. Biome data approximates the majority of occurrences for a species across three ‘super biomes’ (temperate/tropical/arid), which were classified based on the terrestrial ecoregions and biomes of Olson et al. (2001).

We assembled seven dispersal-related characters to assign species to dispersal syndromes: fruit dehiscence, fruit fleshiness, fruit size, seed size, diaspore release, the presence of additional specialized dispersal adaptations (e.g., wings), and the identity of the dispersed unit (i.e., seed, whole fruit, fruit with attachments [e.g., persistent floral organs, such as bracts], seeds with specialised covering [e.g., aril]) (detailed definitions in **Table S1**). In addition, in order to combine knowledge on extant species with discoveries from the fossil record and improve the reconstruction of historical syndromes, ten species from the conservative fossil-calibration set in Ramírez-Barahona et al. (2020) that included recognizable fruits were scored for the same characters as the extant taxa.

We used a trait-based approach, focusing on diaspore structure and morphology, to assign species to six dispersal syndromes: animal (zoochory), wind (anemochory), water (hydrochory), ballistic or self-propelled (autochory *sensu stricto*), gravity (barochory), and prevention or strong limitation of seed dispersal (antitelochory) (**Table S2**). We use autochory *sensu stricto* here for active dispersal by the plant itself, such as explosive dehiscence, whereas the broader analytical category of autochory used below also includes barochory and antitelochory. We acknowledge limitations of syndrome classification (Sinnott-Armstrong et al., 2022), and therefore preferred explicit descriptions of traits in floras or structural studies (n = 892 species). Where such descriptions were unavailable, direct observations of dispersal (n = 183), database entries such as AusTraits (n = 124) (Falster et al., 2021), or interpretations from research-grade iNaturalist photographs (n = 2) were used. Syndromes were scored at species level for 1081 species. For the remaining 120 taxa, the dispersal syndrome of the closest available relative was recorded. Eight of ten fossil taxa allowed syndrome interpretation from preserved recognizable traits. For additional details on syndrome classification, see Supporting methods and results.

### Data quality and potential sampling biases

To evaluate the sampling, quality and biases of our dataset, we compared our syndrome assignments to the much larger but possibly lower quality dataset of Jin et al. (2026a), given that their sampling was opportunistic and heterogenous (Wasowicz & Green, 2026). The Jin et al. dataset includes 35’131 species (297 families, 59 orders) classified in the same four dispersal syndromes (zoochory, anemochory, autochory, hydrochory) used here. To compare the datasets, we harmonized the taxonomy following the World Flora Online plant list (WFO, 2024) and compared family-level taxonomic coverage, identified shared taxa, and calculated the proportion of shared taxa for which four-state dispersal-syndrome assignments differed between datasets. Further, to assess whether our subsampling can represent the dispersal syndromes detected by deeper species sampling within families, we also quantified within-family syndrome variation in the Jin et al. (2026a) dataset using normalized Shannon entropy calculated across the four dispersal syndrome states, with values close to 0 indicating high support for a single state and values closer to 1 indicating more evenly distributed support among states. Low within-family entropy indicates that species within a family largely share a dispersal syndrome, suggesting that subsampling few species within such families should give similar results to deeper sampling.

All analyses were performed in R version 4.3.1 (R Core Team, 2023). The main packages used for analyses were phytools version 1.9.16 (Revell, 2012), ape version 5.8.1 (Paradis & Schliep, 2019), and ade4 version 1.7.22 (Dray & Dufour, 2007).

### Principal component analysis of dispersal-related characters

To visualise our dispersal-related traits into dispersal syndromes, we performed a principal component analysis (PCA) using the seven dispersal-related traits outlined above. The multi-state character ‘diaspore-type’ was split into four simple binary present-absent characters for each state, and proportional seed size (seed size divided by fruit size) was calculated, resulting in a total of eleven characters. Seed size, fruit size, and proportional seed size were log-transformed and the presence-absence of any nutritious structure was used to represent ‘other specialized adaptations’. To perform the PCA, the *dudi.mix* function from the ade4 package was used, as this function is specifically intended for mixed quantitative and discrete data (Dray & Dufour, 2007).

### Ancestral dispersal syndrome reconstructions

To infer the evolution of dispersal syndromes, ancestral state reconstructions (ASR hereafter) using an empirical Bayesian (i.e., maximum marginal likelihood) approach were performed. For this, we used the relaxed calibration phylogenetic tree with the complete fossil calibration set of Ramírez-Barahona et al. (2020), which estimated the crown age of angiosperms at 177.0–218.1 Mya. Fossil species for which dispersal-syndrome information was available were added as zero-length terminal branches at the nodes where these fossils were used for calibration, using the *bind.tip* function in the phytools R package (Revell, 2012).

Dispersal syndromes were reconstructed using a four-state model with zoochory, anemochory, hydrochory, and autochory *sensu lato* (**Table S2**). This grouping retains the key difference between dispersal by animals, wind, or water and dispersal without specialised assistance, while avoiding over-parameterization of sparsely represented and relatively similar syndromes. We inferred marginal ancestral-state probabilities at the nodes of the phylogenetic tree using marginal ancestral character estimation. We compared seven continuous-time Markov models using the sample-size-corrected Akaike information criterion (AICc). Four models were fitted with *fitMk*: equal transition rates (ER), symmetrical transition rates (SYM), all transition rates different (ARD), and a custom model in which only transitions to and from autochory were allowed (CUSTOM). In addition, three hidden-rate Markov models were fitted with phytools’ *fitHRM* function: HiddenER, HiddenSYM, and HiddenARD. These hidden-rate models were included to allow for among-lineage heterogeneity in transition rates that may arise from unobserved covariates of dispersal-syndrome evolution, following Boyko & Beaulieu (2021). Because previous work has shown that diversification rates are not associated with dispersal syndrome evolution (Jin et al., 2026a) and the sampling approach used here was not suitable to test this further, we did not include diversification rate models in the comparison.

The model with the lowest AICc was passed to *ancr* in phytools to calculate the marginal ancestral-state probabilities of each node. To infer the frequency of transitions between dispersal syndromes, the best-fitting model was used to run 1000 stochastic character mappings with the *simmap* function in phytools. In addition, to evaluate the relative frequencies of syndromes through time, we applied phytools’ *ltt* function to a random subset of 100 stochastic character maps.

We also reconstructed the evolution of four dispersal-related characters to evaluate the types of fruits early angiosperms had: fleshiness (present/absent), dehiscence (present/absent), nutritious structure (present/absent), and diaspore type (whole fruit/seed/covered seed/floral fruit). We followed the same general approach as described above for ASR and stochastic character mapping (for more details, see Supporting methods and results).

### Sensitivity of ancestral state reconstructions and transition rate inferences to taxon sampling

To evaluate how our restricted (<1% of angiosperm species) but representative (all angiosperm families represented) sampling could bias ancestral state reconstructions and transition rate inferences, we performed a stratified jackknife analysis. Taxa were removed at five levels: 10%, 20%, 30%, 40%, and 50%, with 1000 (ASR) or 100 (transition rate) replicates per level. In each replicate, approximately equal proportions of taxa were removed from each order, while retaining at least two taxa per order. Orders represented by two or fewer taxa were therefore not pruned. This approach reflects the sampling design of the original dataset, which was assembled specifically to represent all angiosperm orders, families and major subfamilies without including all angiosperm species. The zero-length fossil tips were retained in all replicates. For each replicate, we recalculated ancestral states and transition rates, and calculated the Hellinger distance for each node, which indicates how different the four-state syndrome distributions are compared to the full-data reconstruction. In addition, we recorded whether the highest-probability state (e.g., zoochory) at each retained node was identical between the full-data and jackknife reconstructions. Finally, we evaluated whether older or more ambiguous nodes were more sensitive to taxon removal, by running a linear model with the mean Hellinger distance (or proportion of replicates recovering the same highest-probability node state) as the response variable, and node age and uncertainty (ambiguity) in the original ASR as explanatory variables. Original ASR uncertainty was quantified as normalized Shannon entropy of the full-data marginal state probabilities for each node. For more details on these sensitivity analyses, see SI supporting methods and results.

### Correlated evolution between dispersal, pollination and biome

We assessed whether dispersal syndrome (zoochory/abiotic) and biome (temperate/tropical/arid, data from Ramírez-Barahona et al., 2020) evolution, and dispersal syndrome and pollination mode (biotic/abiotic, data from Stephens et al., 2023) evolution, were correlated, using correlated evolution models (Pagel 1994). In addition, we assessed which dispersal-related characters may have been influential in biome/dispersal syndrome transitions, by assessing correlated evolution between biome or dispersal syndrome and fleshiness or dehiscence. In essence, Pagel’s model works by combining two separate binary characters into a single character, which has the four possible character state combinations as its states. For this combined character, it is then tested whether transition rates between the states of each character differ depending on the state of the other character. As we were interested in identifying whether transition rates in one character were contingent on the state of the other, unidirectional dependent models were constructed, in which the transition rates of only one character depended on the state of the other character. We adapted the model to fit the three-state biome character, by coding a transition matrix with the six possible combined states. For all character combinations, 20 iterations were run using phytools’ *fitMk* function for the interdependent (correlated evolution) model, both ‘adapted’ models with one dependent character, and the independent (non-correlated evolution) model. We used AICc to identify the best-fitting model.

## Results

### Data quality and potential sampling bias

Our dataset comprised 1,201 angiosperm taxa representing 434 families. Across tips, zoochory was the most common syndrome (522 species, 43.5%), followed by anemochory (343 species, 28.6%), autochory (296 species, 24.6%), and hydrochory (40 species, 3.3%).

Comparison with the broader dispersal-syndrome compilation of Jin et al. (2026) showed substantial differences in taxonomic coverage and syndrome interpretation. Our curated dataset included 418 angiosperm families after WFO harmonization, compared with 297 angiosperm families in Jin et al. All angiosperm families represented in Jin et al. were also present in our dataset, whereas 121 families were unique to our dataset, including Amborellaceae. Because Amborellaceae represents the extant sister lineage to all other extant angiosperms, its absence means that the deepest extant angiosperm split was not represented in the Jin et al. analysis. Syndrome assignments were different for 119 of the 391 shared species (30.4%), indicating that the datasets differ not only in taxonomic coverage, but also in syndrome interpretation. Within-family syndrome variation in the Jin et al. dataset was generally low, suggesting that species within a family share a similar syndrome, with a mean normalized Shannon entropy of 0.275 across families (median = 0.201) **(Fig. S1)**.

### Principal component analysis of dispersal-related characters

The principal component analysis (**Fig. 2**) illustrates that the largest physical trait difference in syndromes is between biotic and abiotic seed dispersal. The primary axis explained 38.7% of variation among characters and the second axis 16.3% (for loadings of characters on axes, see **Table S3**). By grouping the species by dispersal syndromes, we found that zoochorous species clustered opposite to anemochorous and autochorous species, illustrating that species with zoochorous dispersal have a pattern of larger seeds, larger fruits, fleshiness, and nutritious structures compared to anemochorous and autonomous species. Anemochorous and autochorous species mostly overlapped, thus showing similarities in dispersal-related characters. Diaspore type (fruit or seed) most clearly distinguished between these two syndromes. Hydrochorous species showed no clear clustering, indicating the lack of unique characters associated with this syndrome. Zoochorous species were split mostly between species where the whole fruit is dispersed, and species with dehiscent fruits, where the dispersal unit is a covered seed (e.g., with aril). Few species are notably separate from the other zoochorous species. These dry non-nutritious species likely represent epizoochorous dispersal syndromes (e.g., *Neurada procumbens, Ibicella lutea*).

**Figure 2:**
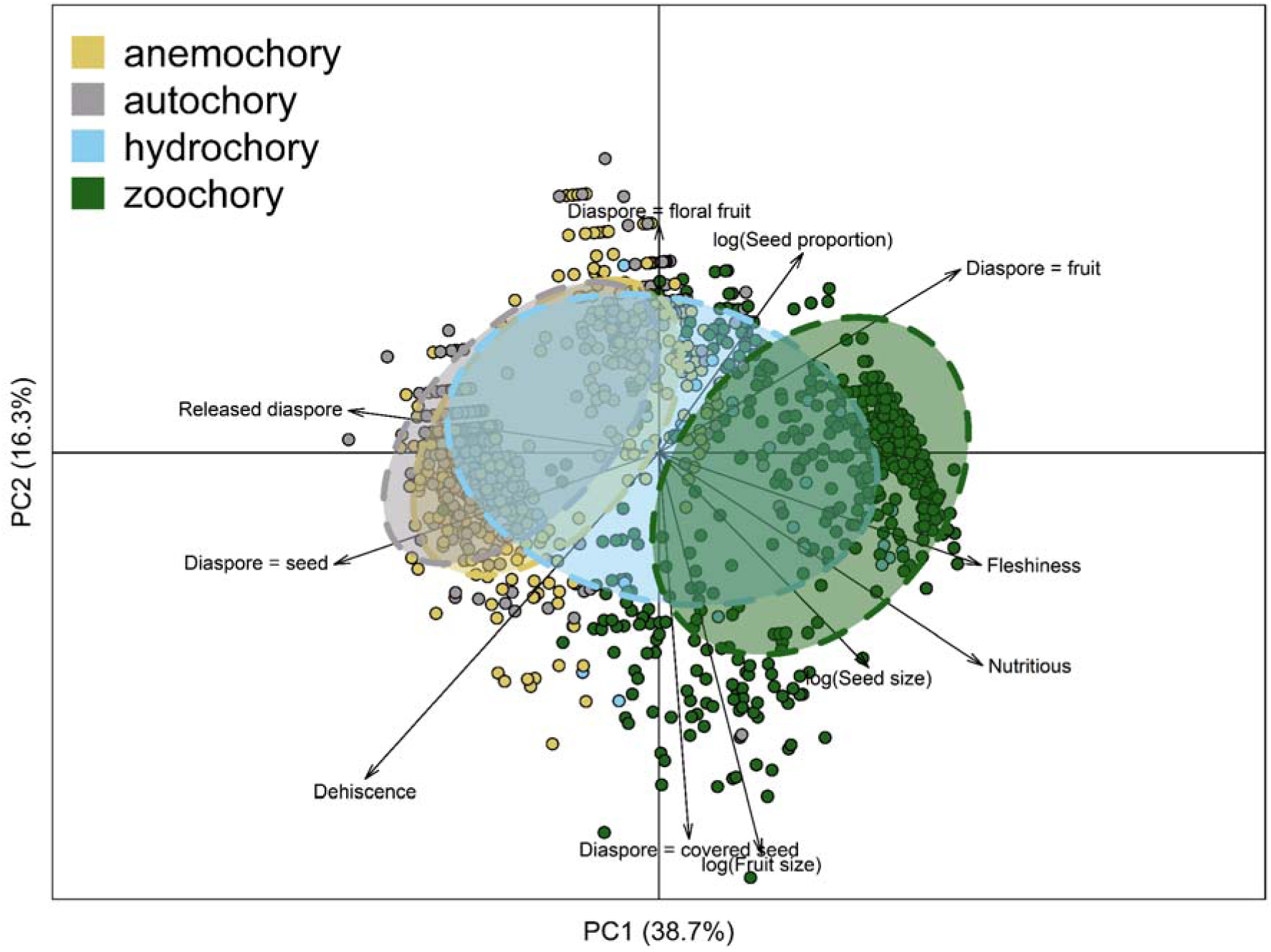
Principal component analysis of dispersal-related characters and associated dispersal syndromes across angiosperms. Points represent species (n = 1201) and are coloured corresponding to their dispersal syndrome. Axes show the relative weight of the corresponding principal component. Arrows represent the loading of the characters analysed. Ellipses were drawn to include 66.7% of species with that syndrome. The arrows with ‘Diaspore = …’ are the four states of the ‘diaspore type’ character (definitions in **Table S1**). ‘Nutritious’ refers to the presence of any nutritious structures.

### Ancestral dispersal syndromes and transition rates

The ancestral dispersal syndrome in angiosperms was found to be zoochory (probability=0.97) (**Figs 3, S2**), likely with indehiscent fruits and whole fruits as the diaspores, but whether these were fleshy or had nutritious structures remains ambiguous (**Table S4**).The ARD model (without hidden states) had the best support (**Table S4**), showing that crown magnoliids (0.99), monocots (0.92), and eudicots (0.80) were ancestrally also most likely zoochorous, whereas rosids (0.98), malvids (0.94), fabids (0.94), asterids (1.0), campanulids (0.99), lamiids (0.98), and commelinids (0.93) were likely autochorous (**Fig. S2**). Of the 59 monophyletic angiosperm orders (>1 taxon) in this dataset, 33 (56%) (25 with probabilities >0.67) were reconstructed to be ancestrally autochorous, 20 (34%) (16 with probabilities >0.67) as zoochorous, six (10%) (four with probabilities >0.67) as anemochorous, and none as hydrochorous (**Figs 3, S2**).

**Figure 3:**
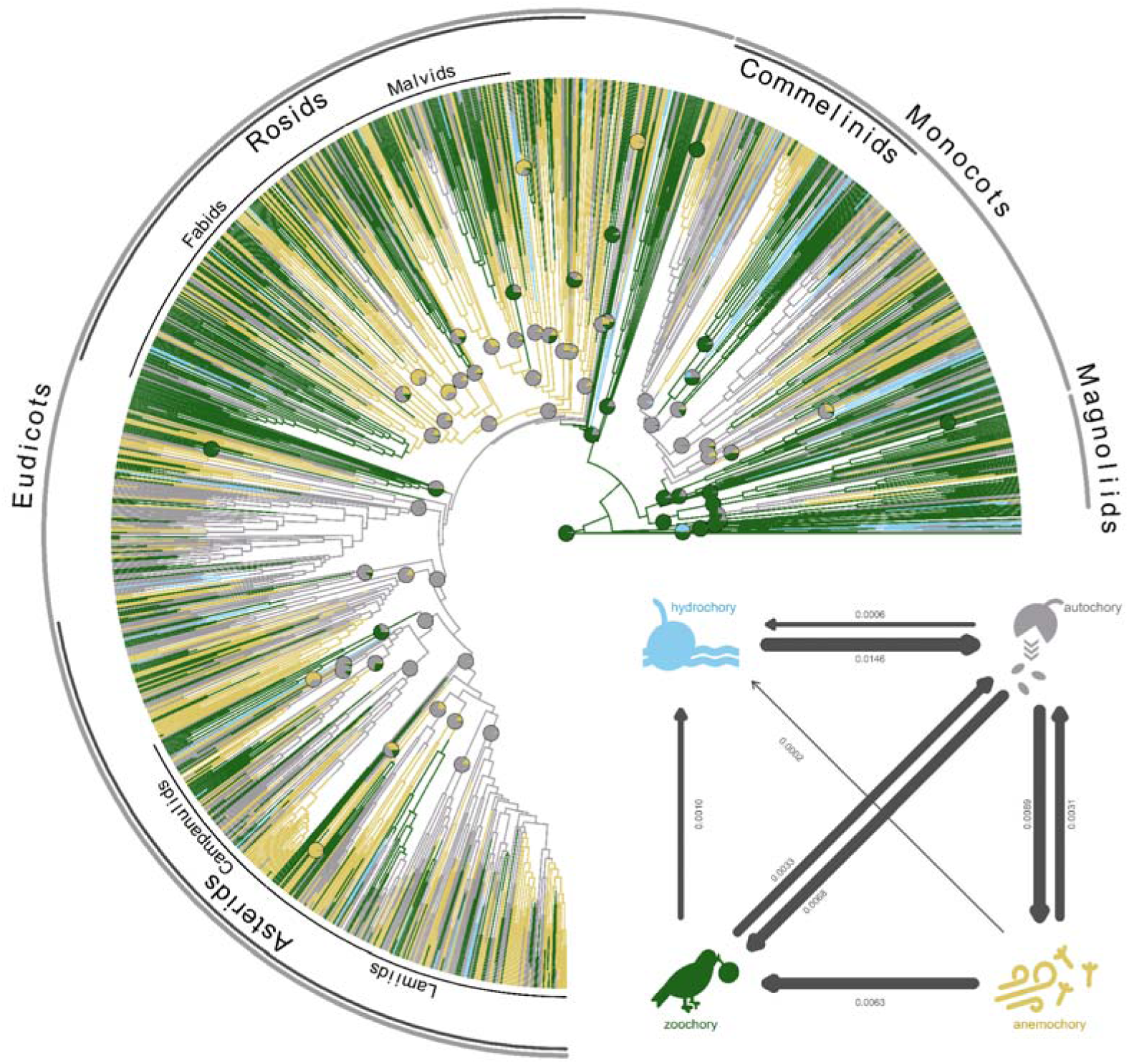
Dispersal syndrome evolution across angiosperms. Ancestral state reconstruction of dispersal syndromes (anemochory [wind]; autochory [barochory/autochory/antitolochory]; hydrochory [water]; zoochory [animal]) across angiosperms under the all rates different (ARD) transition rate model. The phylogeny includes 1201 species, with representatives of all angiosperm families. Pie charts represent marginal ancestral likelihoods and are indicated at crown nodes of orders and major clades as named on the outside perimeter. Colours along branches represent dispersal syndromes as reconstructed in one randomly selected stochastic character map. Transition rates are expressed in number of transitions per million years, with arrow thickness proportional to the rate. Missing transition rate arrows are <0.00005 lineages Myr^-1^.

The highest transition rate (**Fig. 3**) was found from hydrochory to autochory (0.0146 transitions Myr^-1^) and the lowest rates from zoochory (0.0033 transitions Myr^-1^) or anemochory (0.0031 transitions Myr^-1^) to autochory. Hydrochory stands out because transition rates into this syndrome were low (<0.0011 transitions Myr^-1^). Furthermore, the transition rates from zoochory to anemochory, and from hydrochory to anemochory or zoochory, were ∼0, indicating that the evolution of anemochory almost always required autochory as the intermediate state. On average, based on the 1000 stochastic character map simulations, ∼786 transitions (95% highest posterior density interval 738-833) between dispersal syndromes occurred throughout angiosperm evolutionary history, given our species sampling (**Table S5**). Lineage-through-time analyses showed multiple changes in dispersal-related traits from approximately 150 Mya onward, contemporaneous with changes in the dispersal syndromes (**Fig. S3**). Zoochory, indehiscence, and whole fruits as diaspores declined in relative frequency as other dispersal syndromes and diaspore types increased, whereas temporal trends in fruit fleshiness and nutritious structures were less pronounced and more ambiguous.

### Sensitivity of ancestral state and transition rate inferences to taxon sampling

Ancestral-state reconstructions of dispersal syndromes and transition rate inferences were only moderately sensitive to stratified taxon removal. For ancestral states, mean Hellinger distances between the full-data and jackknife reconstructions increased gradually with the proportion of taxa removed, from 0.034 at 10% removal to 0.120 at 50% removal **(Fig. 4a)**. The same pattern was found for the mean percentage of nodes that recovered the same highest probability state between the full and jackknifed data, with 96.5% at 10% removal to 88.0% at 50% removal (**Fig. 4b**). For transition rates, transitions among the three most frequent syndromes (anemochory, zoochory, autochory) remained similar in terms of absolute values across jackknife reconstructions, except for the rate from zoochory to anemochory, which was <0.0001 in the full dataset and on average 0.0013 at 50% taxon removal (**Figs 4c, S4**). Additionally, transition rates related to hydrochory, the rarest state (3.3% extant taxa), were sensitive to taxon removal, with increasingly skewed rate distributions with more taxa removed. This also caused moderate decreases in relative contributions of rates between anemochory, zoochory and autochory (**Figs 4c, S4**). We found that both original ASR uncertainty (i.e., ambiguity in ancestral syndrome state in the full-data reconstruction) and node age increased the mean Hellinger distance between the full-data and jackknife reconstructions (**Fig. S5**), with original ASR uncertainty consistently being the strongest predictor. This suggests that ambiguous nodes remained ambiguous after taxon removal, and these were not necessarily deeper (older) nodes in the phylogeny. In contrast, less ambiguous nodes consistently had syndrome reconstructions with high confidence, even when 50% of taxa were removed.

**Figure 4.**
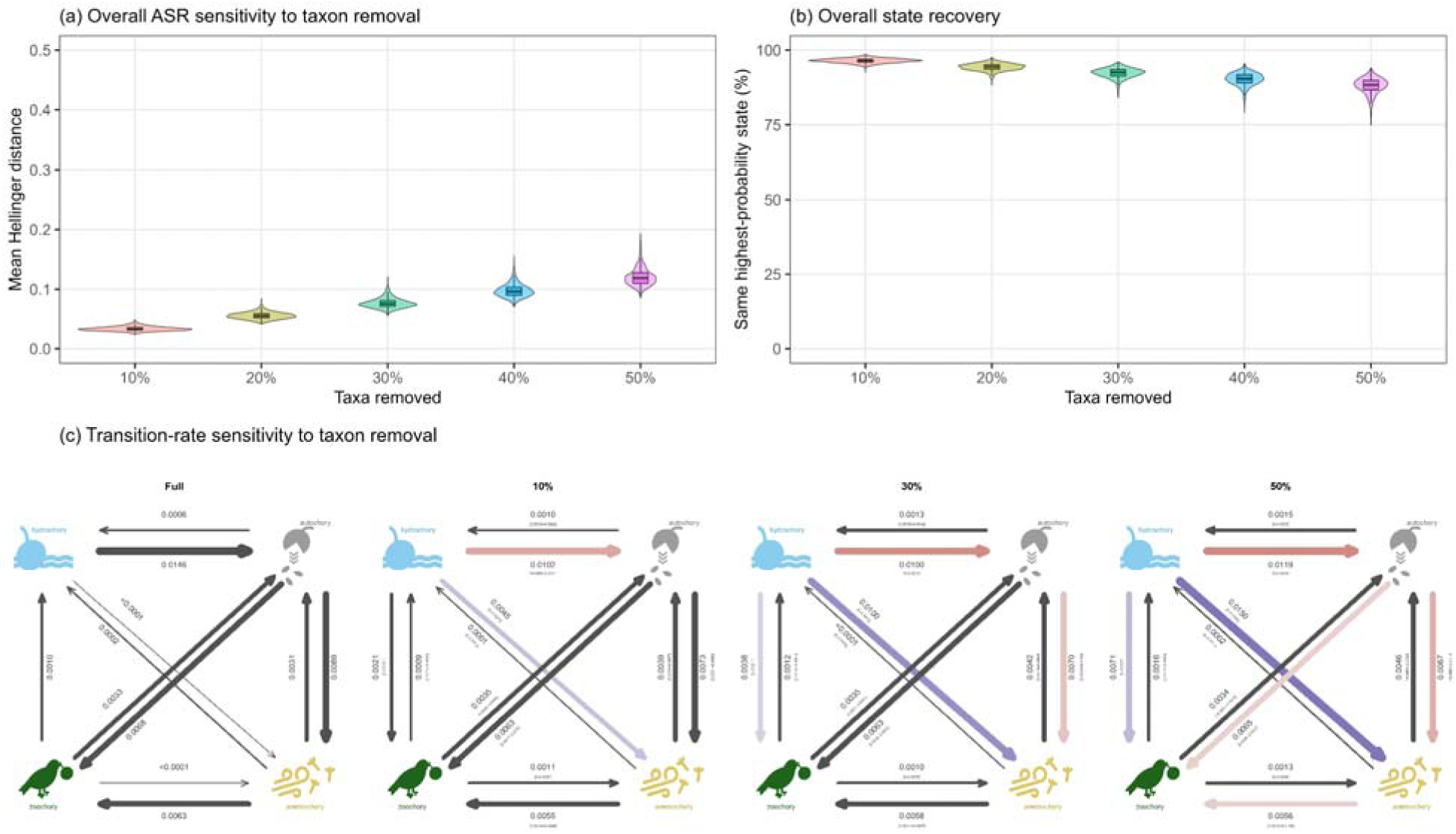
Sensitivity of ancestral-state reconstructions and transition rates to taxon removal. (a) Replicate-level mean sensitivity across taxon-removal levels. (b) Percentage of retained nodes recovering the same highest-probability state as the full-data reconstruction across jackknife replicates at each removal level. (c) Mean ARD transition-rate estimates for the full dataset and after 10%, 30%, and 50% order-stratified taxon removal (for 20% and 40%, see **Fig. S4**); brackets show empirical 2.5–97.5% quantiles across 100 jackknife replicates. Coloured arrows indicate transitions whose mean share of the total rate vector changed by ≥5 percentage points relative to the full dataset, where red represents a decrease and blue represents an increase.

### Correlated evolution between dispersal, pollination and biome

We found support for correlated, rather than independent, evolution for all pairs of characters (**Figs 5, S6, Table S6**). Specifically, transition rates between biomes or pollination modes were dependent on dispersal syndromes, but not *vice versa* (ΔAICc = 32.7 and 2.6 respectively, vs. independence; all model comparisons are provided in **Table S6**). For biome transitions, we found that transition rates from tropical to temperate or arid biomes were at least twice as high for lineages with abiotic than zoochorous dispersal (0.006/0.003 transitions Myr^-1^ with abiotic vs. 0.003/0 with zoochorous dispersal), while transition rates from temperate or arid to tropical biomes were at least twice as high for lineages with zoochorous than abiotic dispersal (0.023/0.006 transitions Myr^-1^ with zoochorous vs. 0.011/0.002 with abiotic dispersal). Similarly, transition rates between temperate and arid biomes were ca. twice as high for lineages with abiotic compared to zoochorous dispersal (**Fig. 5a**). For pollination mode transitions, we found that transition rates from abiotic to biotic pollination were dependent on zoochorous dispersal (0.003 transitions Myr^-1^ with zoochorous dispersal vs. ∼0 with abiotic dispersal), whereas transitions from biotic to abiotic pollination depended on abiotic dispersal (0.001 transitions Myr^-1^ with abiotic dispersal vs. ∼0 with zoochorous dispersal) (**Fig. 5b**). In contrast, transition rates between dispersal syndromes were not conditional on biome or pollination mode, but they were influenced by the evolution of fleshiness or dehiscence (**Fig. S6**). Specifically, transitions to zoochorous dispersal were higher in lineages with fleshy or dehiscent fruits, and transitions to abiotic dispersal higher in lineages with dry fruits.

**Figure 5:**
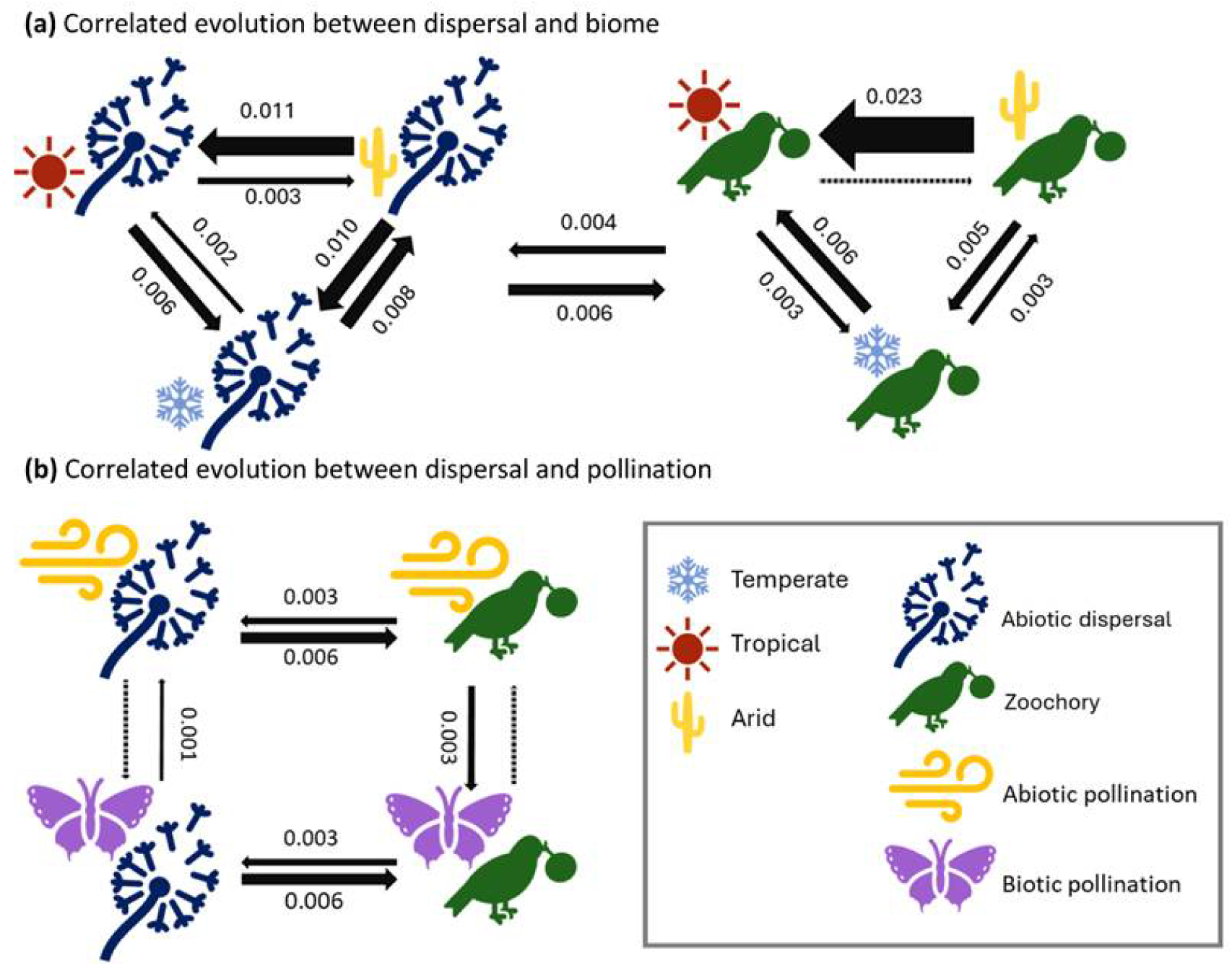
Correlated evolution between dispersal syndrome, biome, and pollination mode evolution in angiosperms. Transition rates based on the best-fitting models for correlated evolution, with arrow width scaled to transition rates. (a) rates of biome shifts (temperate/tropical/arid) are dependent on dispersal syndrome (zoochory/abiotic). The arrows in the middle represent transitions between dispersal syndromes for any of the biomes, as transitions between dispersal syndromes were inferred to be independent from biome shifts. (b) evolution of pollination mode (abiotic/biotic) is dependent on dispersal syndrome (zoochory/abiotic). Rates are presented in transitions Myr^-1^. Dashed arrows indicate a transition rate <0.0005 transitions Myr^-1^.

## Discussion

We inferred the evolutionary history of seed dispersal mechanisms in angiosperms using a novel high quality curated dispersal-related character and syndrome dataset (**Figs 2, 3, Table S2**), which included representative species of all angiosperm families. Ancestral state reconstructions showed that the ancestor of all angiosperms, multiple major clades (magnoliids, monocots, and eudicots), and 20 orders were most likely dispersed by animals (zoochory). In contrast, most orders (33) had autochorous dispersal (**Figs 3, S2)**. Our reconstructions were mostly robust against taxon removal scenarios (**Figs 4, S5**) and are consistent with inferences made for clades with much denser species sampling (e.g., anemochory in Fagales, Larson-Johnson 2016), illustrating that our sampling of >1200 angiosperm species across representatives of all extant angiosperm families can provide reasonable inferences, at least for early angiosperm evolution. In support of H1, we found that transitions between dispersal syndromes have occurred frequently. For example, dispersal syndrome transitions happened five times more frequently than pollination mode transitions based on the same phylogeny and species sampling (∼786 vs. ∼146 transitions, Stephens et al., 2023). Nevertheless, transition rates were similar across dispersal syndrome states, except for the apparent absence of transitions from zoochory to anemochory, which required a stepwise transition via autochory. Additionally, in support of H2, we detected correlated evolution between dispersal syndromes and biomes, as well as between dispersal syndromes and pollination modes (**Fig. 5, Table S6**). This suggests that transitions between zoochory and abiotic dispersal may have predisposed lineages to shift between biomes or pollination modes, critical for the ecological diversity of angiosperms.

### The most recent common ancestor of angiosperms had fruits dispersed by animals

We show that the most recent common ancestor of angiosperms probably relied on zoochorous seed dispersal, with fleshy, indehiscent fruits, which resolves uncertainty in previous angiosperm-wide assessments, which indicated zoochory and/or autochory as the ancestral state (Jin et al., 2026a), and earlier hypotheses based on paleontological data (Tiffney, 1984; Tiffney, 2004; Wing & Tiffney, 1987; Eriksson et al., 2000). However, our findings are consistent with recent discoveries of large diaspores from an in situ late Campanian (74.6 million years ago) tropical forest from the Jose Creek Formation in New Mexico (Lee et al., 2026). These early angiosperms may have been understory bushes in moist tropical environments (Kerkhoff et al., 2014; Pouteau et al., 2020), with insect-pollinated flowers (Stephens et al., 2023), woody stems (Philippe et al., 2008), and with a superior gynoecium with more than five apocarpous carpels (Sauquet et al., 2017; Xiang et al., 2024). Interestingly, today, zoochorous angiosperms in the wet tropics are most frequently dispersed by birds and mammals (Correa et al., 2015). However, true birds only emerged ∼67 Mya (Stiller et al., 2024) and even when including extinct relatives, they originated between ∼165 and 150 Mya (Brusatte et al., 2015). Although mammals originated between ∼251 and 165 Mya, their rapid diversification started only about 66 Mya (Álvarez-Carretero et al., 2022). As the estimated crown age of angiosperms predates these ages (Sauquet et al., 2022), including the one used here (∼197 Mya), this raises the question of which animals were responsible for seed dispersal in ancestral angiosperms. Herbivorous lizards, dinosaurs, or early synapsids were already widespread at this time and could thus conceivably have been dispersers for early angiosperms, but convincing evidence from coprolites, for example, is missing (Tiffney, 2004, but see Rodrı guez-de la Rosa et al., 1998). Further support for zoochory as the ancestral condition is found in the presence of different conserved groups of genes that are involved in fruit development and ripening, which have been found in angiosperms and gymnosperms, implying that the molecular machinery required to produce fleshy fruits was likely already present in ancient angiosperms (Lovisetto et al., 2012).

### Transitions and constraints in dispersal syndrome evolution

We detected frequent transitions between dispersal syndromes throughout angiosperm evolution, at least at the phylogenetic scale examined here. Although zoochory was likely abundant in early angiosperms, the lineage-through-time analyses suggest that a broader diversification of dispersal strategies began around 150 Mya: zoochory, indehiscence, and dispersal of whole fruits, which were initially dominant, declined in relative frequency as other dispersal syndromes and diaspore types became increasingly common (**Fig. S3**). Changes in diaspore characters, such as dehiscence and diaspore type, may have been critical adaptations that facilitated the evolutionary transition to different dispersal vectors and syndromes, consistent with results from correlated evolution models (**Fig. S6**). Patterns for fleshiness and nutritious structures were less pronounced, however, and their early reconstructions were comparatively uncertain (**Fig. S3**). This diversification of dispersal adaptations, together with the strong homoplasy of syndromes in relation to the characters involved (**Fig. 1**) (Wake et al., 2011) and frequent transitions among dispersal syndromes, may in turn have facilitated the widespread dispersal and diversification of angiosperms during the ATR (Benton et al., 2022). Dispersal syndrome evolution in angiosperms thereby illustrates how the repeated reinvention of traits and trait combinations may be associated with the ecological and evolutionary diversification of angiosperms (i.e., ‘Darwin’s abominable mystery’) (Crepet & Niklas, 2009).

Nevertheless, we also detected evolutionary constraints in dispersal syndrome evolution. For example, transitions from zoochory to anemochory and from hydrochory to zoochory or anemochory have been entirely absent. Instead, autochorous dispersal seemingly served as an intermediate step. This points to a stepwise transition between certain dispersal syndromes. To illustrate: monocots were reconstructed as zoochorous, but there is a shift to autochorous dispersal after the split with Acorales and Alismatales. After this, several shifts to anemochory were reconstructed, such as the one in Poales, at the base of Bromeliaceae (**Fig. S2**). Such a transition could occur via stepwise trait changes, because transitions in the traits that make up syndromes inherently cause transitions between syndromes. For example, a fleshy indehiscent zoochorous species first becomes barochorous or autochorous by losing or modifying fleshy structures, before developing wing-like structures that facilitate more specialised dispersal through anemochory. In contrast, the reverse transition from anemochory to zoochory has occurred frequently (∼129 transitions, **Table S5**), possibly because a one-step transition is possible. For example, from a wind- to epizoochorous fruit (e.g., in Asteraceae), or from a dehiscent fruit with dust-like seeds to an indehiscent fruit enabling endozoochory, without the need to lose the dust-like seeds first (e.g., *Vaccinium uliginosum*). Stepwise evolution of complex characters, such as from zoochory to anemochory, has previously been implied for roots (Hetherington & Dolan, 2018) and pollination modes (Culley et al., 2002; Stephens et al., 2023). For example, the evolution of roots in lycopsids first required the development of a positively gravitropic non-leafy shoot, followed by the evolution of a root cap, root hairs, and more (Hetherington & Dolan, 2018). This suggests that the evolution of complex syndromes may often involve stepwise trait evolution.

### Dispersal syndromes facilitated transitions between biomes and pollination modes

We show that dispersal syndrome evolution may have affected shifts between major biomes and pollination modes. Specifically, we found that zoochorous dispersal was associated with increased transition rates to tropical biomes (from arid or temperate biomes), while abiotic dispersal was associated with increased transition rates to arid or temperate biomes (from tropical biomes), or transitions between arid and temperate biomes (**Fig. 5a**). This finding suggests that the evolution of zoochory, which is advantageous in closed tropical forests that host a wide diversity of seed dispersing animals (Vasconcelos et al., 2023), and abiotic dispersal, which is advantageous in more open biomes (Correa et al., 2015), may have preceded the transition to the biome for which the syndrome is advantageous. This fits with a ‘trait first’ rather than a ‘biome first’ model of evolution for dispersal syndrome (Zanne et al., 2014). For example, in Torricelliaceae the shift from autochorous to zoochorous dispersal appears to predate its reconstructed transition from temperate to tropical biomes (Ramírez-Barahona et al., 2020). Furthermore, this environmental match may affect speciation: fleshy-fruited zoochorous species have higher speciation rates in closed canopy tropical forests than dry-fruited non-zoochorous species, but zoochorous species that inhabit more open habitats show lower speciation rates (Givnish, 2010). Transitions in dispersal syndromes may therefore have predisposed lineages to radiate in new biomes, rather than arising as adaptations after biome shifts. Consistent with this, diversification appears to be less associated with the syndrome itself (Jin et al., 2026a) than with its interaction with the environment (Donoghue & Sanderson, 2015; Bouchenak-Khelladi et al., 2015; Onstein et al., 2017).

We also found that transitions between biotic and abiotic pollination were dependent on dispersal syndrome, with transitions to biotic pollination primarily occurring in zoochorous species, and *vice versa* for abiotic dispersal (**Fig. 5b**). This suggests that pollination and seed dispersal roles may be evolutionarily linked (e.g., as in ‘double mutualists’), and seed dispersal by animals may have favoured evolutionary transitions back to animal-mediated pollination. Support for such dependent evolution was also found in lizards, where a fruit-eating diet preceded their ability to also pollinate flowers (Kahnt et al., 2023) and across plants between their mutualism types, including pollination, seed dispersal, and mycorrhizal fungi (Yamawo & Ohno, 2024). Environmental selection may explain the association between pollination and seed dispersal syndrome evolution. For example, plants living in dry, cold or open, high latitude environments with few available animals may have higher rates of abiotic pollination and seed dispersal, and *vice versa* (Chen et al., 2017, Stephens et al. 2023). In summary, we conclude that dispersal syndrome evolution in angiosperms is closely associated with the abiotic (e.g., biome) and biotic (e.g., pollination, frugivory) environment.

### Sampling effects and limitations

By using a trait-based approach to assign species to dispersal syndromes, we addressed recent criticism on syndrome assignments (Wasowicz & Green, 2026) of a more broadly-sampled dataset (Jin et al., 2026a). However, although our taxon removal approach illustrates limited effects of sampling inferred ancestral syndrome states and transition rates (**Fig. 4**), our inferences of transition rates show differences of about an order of magnitude with rates inferred in Jin et al., (2026a). This suggests that our rates are likely underestimated due to long terminal branches (representing families), whereas they may have been overestimated in Jin et al., (2026a) due to both data error noise (more variation among tips than real) and the way polytomies were artificially resolved in the phylogenetic tree. Additionally, it is important to note that our inferences were based on a molecular phylogeny with limited information from fossil taxa, limited sampling, and simple Mk models that assume constant transition rates through time (but see Reyes et al. (2018)), and our analyses did not include any gymnosperm outgroups, which could improve reconstructions. Expanding our work with more seed and fruit characters from fossil taxa (e.g., as done in López-Martínez et al., 2024 with fossil flowers) and increasing the sampling of extant and extinct species (Leslie et al., 2013; Nigris et al., 2021) could therefore provide additional insights on dispersal syndrome evolution in angiosperms.

## Conclusion

We show that ancestral angiosperms had most likely zoochorous seed dispersal, and we propose that variability in angiosperm diaspore characters enabled the frequent and stepwise evolution of dispersal syndromes in angiosperms, with autochory as an evolutionary step between zoochory and anemochory. Furthermore, dispersal syndrome change has served as an important pre-adaptation to biome and pollination mode shifts, key components of the floristic and ecological diversity of angiosperms (Crepet & Niklas, 2009, Ding et al. 2025). Nevertheless, many open questions remain. For example, which animals were eating and dispersing the ancestral angiosperm fruits, and was that an innovation that arose along the stem lineage of angiosperms, or an older syndrome inherited from deep time seed plant ancestors?

## AI use declaration

ChatGPT (OpenAI; accessed 2025–2026) was used to support manuscript editing, wording of selected sections, and troubleshooting, cleaning, and improving the efficiency of R code originally written by the authors. All AI-assisted text and code were critically reviewed, revised, and verified by the authors, who take full responsibility for the analyses, interpretation, and conclusions.

## Acknowledgements

We thank the members of the Tropical Botany and Biodiversity Hotspots groups and Rutger Vos at Naturalis, and the Evolution & Adaptation research group at the German Centre for Integrative Biodiversity Research (iDiv) Halle – Jena - Leipzig, for discussions and advice. We thank Liam J. Revell and Luke J. Harmon for important insights in phylogenetic comparative methods and instruction and help with phytools.

## Author contributions

REO and DdF designed the research; DdF collected data, performed analyses and prepared figures, DdF and REO wrote a first version of the manuscript, and FTB, HS and RES provided comments and suggestions that led to the final version. REO supervised the study.

## Data Accessibility Statement

All data files, including dispersal-related characters for extant and extinct taxa, and R scripts, are available from Dryad (DOI: TBD). Reviewer URL: TBD.

