## Supporting information for "Dispersal syndrome evolution in angiosperms: ancestral zoochory, correlations with pollination mode and biome, and sampling effects"

Supporting methods and results

### Additional details on syndrome assignments

Some species show characters that could be interpreted for more than one dispersal syndrome. In these cases, we selected the syndrome for which the clearest adaptations were present. For example, *Tragopogon dubius* has characters associated with both anemochory (feathery pappus) and zoochory (hooked appendages) (Sukhorukov & Nilova, 2015). We recorded this species as 'anemochory', because the hooked appendages are small and not openly exposed to allow for (epi-)zoochory. In rare instances (n = 3), species were found to have polymorphic diaspores. In such cases, the syndrome generally responsible for the largest dispersal distance (following Tamme et al., 2014) was recorded. Hydrochorous dispersal was frequently mentioned for species with small light-weight seeds, though these often showed no clear adaptation for prolonged flotation, thus barochory was recorded, as proximity to water and buoyancy alone do not constitute a convincing adaptation for hydrochory.

Interestingly, some angiosperm species have multiple traits that are functional adaptations to different dispersal syndromes. For example, *Buxus sempervirens* (Buxaceae) has explosive dehiscent fruits indicative of autochorous dispersal, but seeds also possess elaiosomes, a clear adaptation for myrmecochory (**Table S2**). Similarly, several species have dimorphic diaspores. For example, fruits of *Aethionema saxatile* (Brassicaceae) can be indehiscent and antitelochorous, or dehiscent and anemochorous, depending on pressure from herbivory (Bhattacharya et al., 2019). We classified the dispersal syndrome of polymorphic species based on the syndrome associated with the furthest dispersal distance (i.e., myrmecochory for *B. sempervirens* and anemochory for *A. saxtatile*), as we considered this the most critical outcome for adaptive evolution in the context of dispersal (Tamme et al., 2014). However, polymorphic dispersal syndromes themselves could be adaptive, and it is unknown how common this is across angiosperms. Additionally, plant species may also disperse vegetatively, such as in *Bambusa vulgaris* (Poaceae), which rarely, if ever, sets seed, but has structures suitable for vegetative reproduction. Finally, some characters may be functional in different dispersal strategies, such as dry or spongy pulp attracting animals (zoochory) as well as facilitating flotation (hydrochory). Such variation could affect our analyses, but comparable cases are probably not sufficiently common to exhibit significant influence on the outcomes and conclusions. Generally, cases like this are illustrative of the flexibility in angiosperm dispersal syndromes and the adaptations used to develop a dispersal strategy, while also recognizing the lack of unique adaptations in certain syndromes (e.g., hydrochory, **Fig. 2 main text**).

### Additional details on ancestral state reconstructions

For all models, we used pi = "fitzjohn" as the root prior, following FitzJohn et al. (2009). To reduce the risk of ending up in local optima, we ran 30 independent random-start optimizations for each conventional Mk model. For the hidden-rate models, we ran 30 independent optimizations with two internal optimization attempts each, resulting in 60 optimization attempts per model. For each model, we retained the optimization with the highest log-likelihood, provided that *fitMk* or *fitHRM* reported convergence and the transition rate values were reasonable (<1).

We reconstructed four additional dispersal-related traits: fruit fleshiness (dry/fleshy), fruit dehiscence (dehiscent/indehiscent), presence of nutritious structures (absent/present), and diaspore type (fruit, seed, covered seed, or floral fruit). Analyses otherwise followed the approach used for dispersal syndrome, including the FitzJohn root prior, repeated random-start optimizations, model comparison using AICc, and calculation of marginal ancestral-state probabilities with ancr. Fossil observations were incorporated as zero-length tips where data were available.

For the three binary traits, the final candidate set comprised ER and ARD Mk models and corresponding two-category hidden-rate models (HiddenER and HiddenARD). No biologically motivated CUSTOM model analogous to that used for dispersal syndrome was included. For the four-state diaspore-type character, ER, SYM and ARD models and their corresponding two-category hidden-rate models were compared. Conventional models were fitted from 20 random starts. Hidden-rate models were fitted with two latent rate categories; the binary HiddenER models used one internal optimization attempt and HiddenARD two, whereas the diaspore hidden models used two internal attempts

For each character, stochastic character maps were simulated under its best-supported evolutionary model. To visualize changes in character-state frequencies through time, 100 stochastic maps were randomly selected and analysed using the *ltt* function in phytools. Lineage counts were converted to relative frequencies at each time point. For traits reconstructed with hidden-rate models, latent categories belonging to the same observed character state were combined before plotting so that the LTT curves represented the biological states rather than hidden rate classes.

### Additional details on sensitivity analyses (taxon removal scenarios)

Modification to approach. For the transition rate inferences, rather than fitting 30 independent optimizations to each replicate as done in the full analysis, we accepted the first run that reported convergence with reasonable rates (<1).

Matching equivalent nodes after taxon removal. To ensure that ancestral-state reconstructions were compared only between equivalent branching events, each internal node of the full tree was defined by its two immediate descendant lineages. After taxon removal, a node was considered retained only if at least one taxon remained in both descendant lineages. If one lineage was lost entirely, the original node was treated as absent and excluded from that replicate. For retained nodes, the corresponding node in the pruned tree was identified as the most recent common ancestor of the remaining descendants of the original node. This procedure prevented comparisons between the original node and a deeper or shallower node created by pruning. Nodes represented in fewer than 5% of replicates were excluded from the final calculations.

Predictors of reconstruction sensitivity. We examined whether sensitivity to taxon removal was associated with properties of the original node.

For each reduced dataset, the phylogeny and corresponding tip-state vector were pruned, and ancestral states and transition rates were recalculated with phytools' ancr (ancestral states) and fitMk (transition rates) respectively. For the ancestral states we used the best-fitting transition-rate model inferred from the full dataset. For the ancestral state inferences, we identified nodes in the taxon-removal scenario that were equivalent to the original node, and compared the jackknife reconstruction with the full-data reconstruction using the Hellinger distance, which is a measure of dissimilarity between two probability distributions, calculated from their marginal ancestral-state probability distributions (Garba et al., 2018). In this case, it calculates how different the four-state syndrome distributions in the nodes are. In addition, we recorded whether the highest-probability state (e.g., zoochory) at each retained node was identical between the full-data and jackknife reconstructions. Results were summarized as the mean Hellinger distance and the percentage of replicates recovering the same highest-probability state for each node and removal level.

To evaluate which nodes were most sensitive to taxon removal, we tested whether the mean Hellinger distance or proportion of replicates recovering the same highest-probability node state were associated with node age and uncertainty (ambiguity) in the original ASR (i.e., syndrome state reconstruction at that node). Node ages were taken from the original dated tree. Original ASR uncertainty was quantified as normalized Shannon entropy of the full-data marginal state probabilities for each node. For each jackknife replicate within each removal level, we fitted a linear model with Hellinger distance as the response variable and with both node age and original ASR entropy included as predictors. The response and both predictors were standardized to a mean of 0 and standard deviation of 1 before model fitting, allowing the resulting regression coefficients to be compared directly. Coefficient distributions were summarized across the 1,000 replicates at each removal level.

Figure S1: Within-family variation in dispersal syndromes in Jin et al. (2026).

Normalized Shannon entropy across the four dispersal syndromes for each WFO-harmonized family; values near 0 indicate dominance by a single syndrome and higher values indicate greater within-family variation.

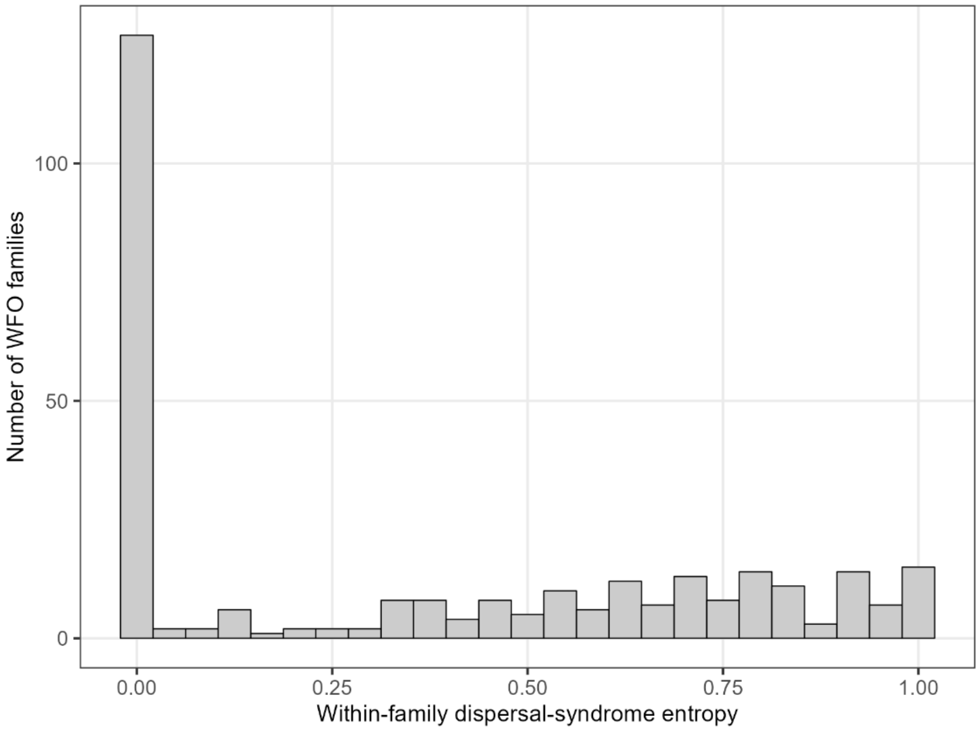

### Figure S2: Dispersal syndrome evolution across angiosperms extended figure.

Marginal ancestral character estimation of dispersal syndromes across angiosperms under the all rates different (ARD) transition matrix. Pie charts represent marginal ancestral likelihoods and are indicated at all nodes. Major clades and orders are named on the right side. Colours along branches represent dispersal syndromes as reconstructed in one randomly selected stochastic character map.

### Figure S3: Relative number of lineages through time for dispersal syndromes, dehiscence, fleshiness, nutritious structures, and diaspore type.

The relative number of lineages through time for dispersal syndrome (a), dehiscence (b), fleshiness (c), nutritious structures (d), and diaspore type (e). Thick lines represent the average proportion of lineages with a trait based on 100 randomly selected stochastic character maps, with thin lines showing individual mappings and associated uncertainty. Hidden states were collapsed with their corresponding observed state. Zoochory, indehiscence, and whole fruits as diaspores were initially dominant and declined in relative frequency from approximately 150 Mya onward, coinciding with increases in other dispersal syndromes, diaspore types, and dispersal-related traits. Temporal patterns in fleshiness and nutritious structures were less pronounced and showed greater uncertainty in the early reconstructions. Due to the low number of lineages before ~150 Mya, patterns earlier than this provide limited information.

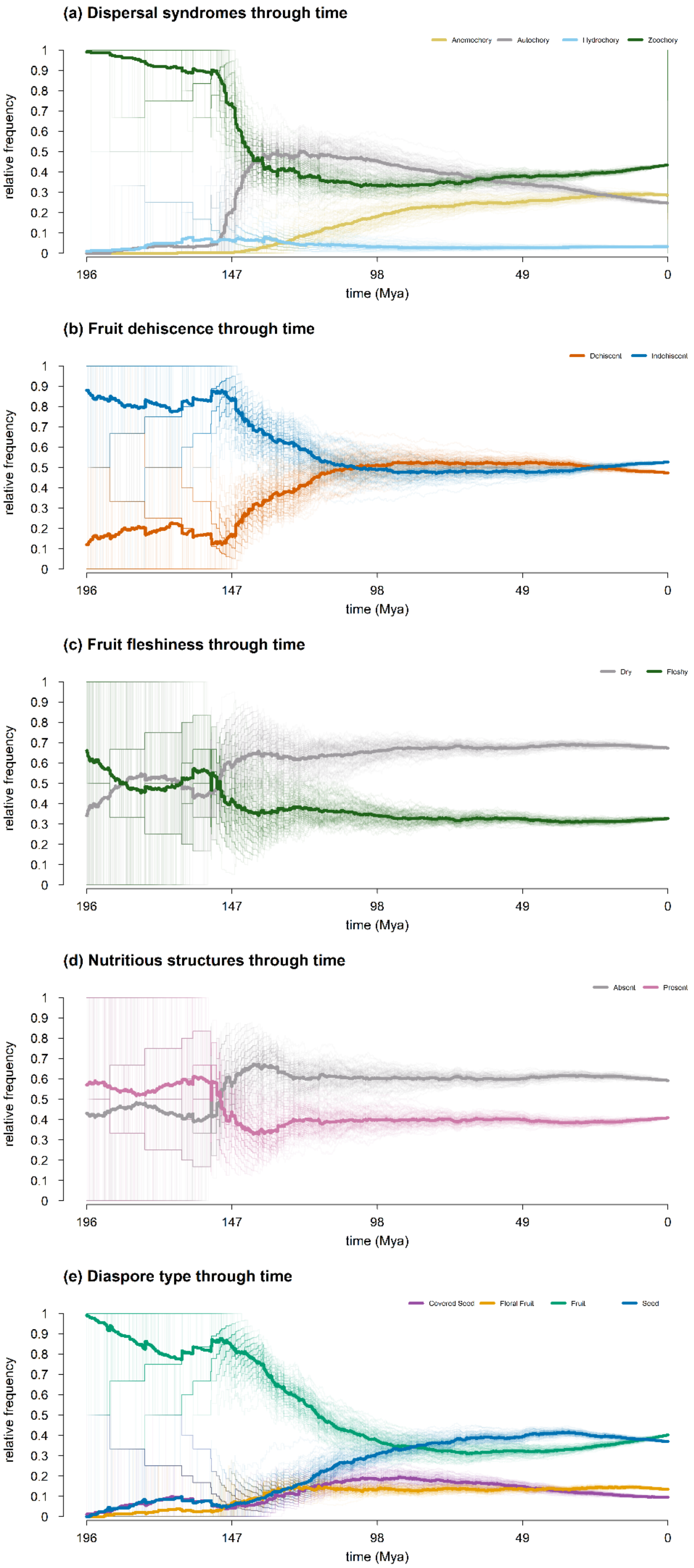
(a) Dispersal syndromes through time
Anemochory
Autochory
Hydrochory
Zoochory
relative frequency
time (Mya)
(b) Fruit dehiscence through time
Dehiscent
Indehiscent
relative frequency
time (Mya)
(c) Fruit fleshiness through time
Dry
Fleshy
relative frequency
time (Mya)
(d) Nutritious structures through time
Absent
Present
relative frequency
time (Mya)
(e) Diaspore type through time
Covered Seed
Floral Fruit
Fruit
Seed
relative frequency
time (Mya)

Figure S4: Sensitivity of dispersal-syndrome transition-rate estimates. to taxon removal.

(a) Transition-rate networks for the full dataset and for order-stratified removal of 10–50% of taxa. For each removal level, values show the mean transition rate across 100 jackknife replicates, with empirical 2.5–97.5% quantiles in brackets; arrow width is proportional to the absolute rate. Coloured arrows indicate transitions whose mean contribution to the total rate vector changed by ≥5 percentage points relative to the full dataset. (b) Percentage-point change in each transition's share of the summed mean transition-rate vector relative to the full-data estimate; purple indicates an increased share and red a decreased share.

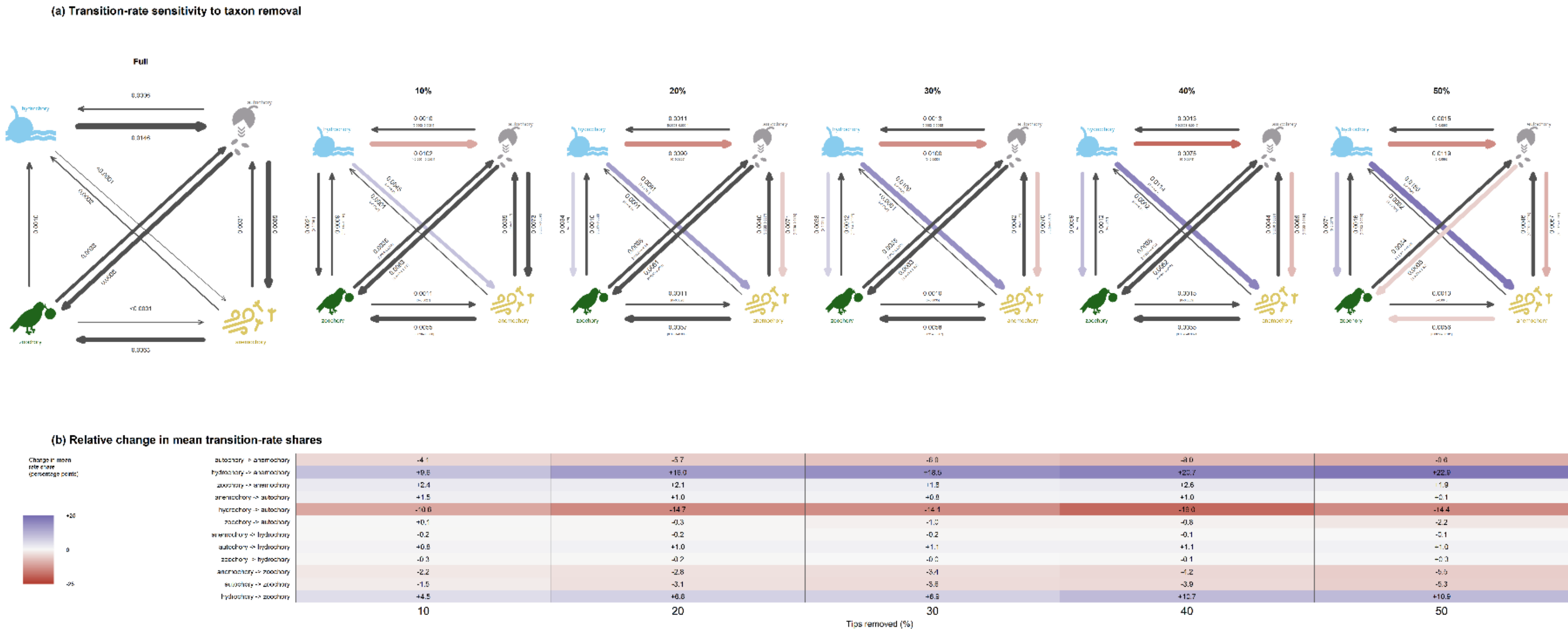

### Figure S5: Effects of original ASR uncertainty and node age on reconstruction sensitivity to taxon removal.

(a–b) Mean Hellinger distance and recovery of the same highest-probability state across quantiles of original ASR uncertainty, grouped by taxon-removal level. Entropy quantiles: Q1 = 0–0.08, Q2 = 0.08–0.22, Q3 = 0.22–0.41, Q4 = 0.41–0.55, and Q5 = 0.55–0.94. At 50% taxon removal, at least 95% of nodes in Q1–Q3 retained the same highest-probability state as in the full-data reconstruction, compared with 86% in Q4 and 78% in Q5. (c–d) Equivalent analyses across node-age quantiles: Q1 = 3.1–33.0 Mya, Q2 = 33.0–51.1 Mya, Q3 = 51.1–70.0 Mya, Q4 = 70.0–97.8 Mya, and Q5 = 97.8–195.8 Mya; the association with node age was weaker than that with original ASR uncertainty. (e) Standardized effects of original ASR uncertainty and node age on Hellinger distance from replicate-level multiple regressions including both predictors. Median standardized effects of original ASR uncertainty ranged from 0.25 to 0.30 across removal levels, meaning that a 1-SD increase in uncertainty was associated with a 0.25–0.30-SD increase in Hellinger distance while controlling for node age. Corresponding effects of node age were smaller, ranging from 0.06 to 0.14 SD while controlling for ASR uncertainty. Boxplots show the distribution of standardized coefficients across jackknife replicates.

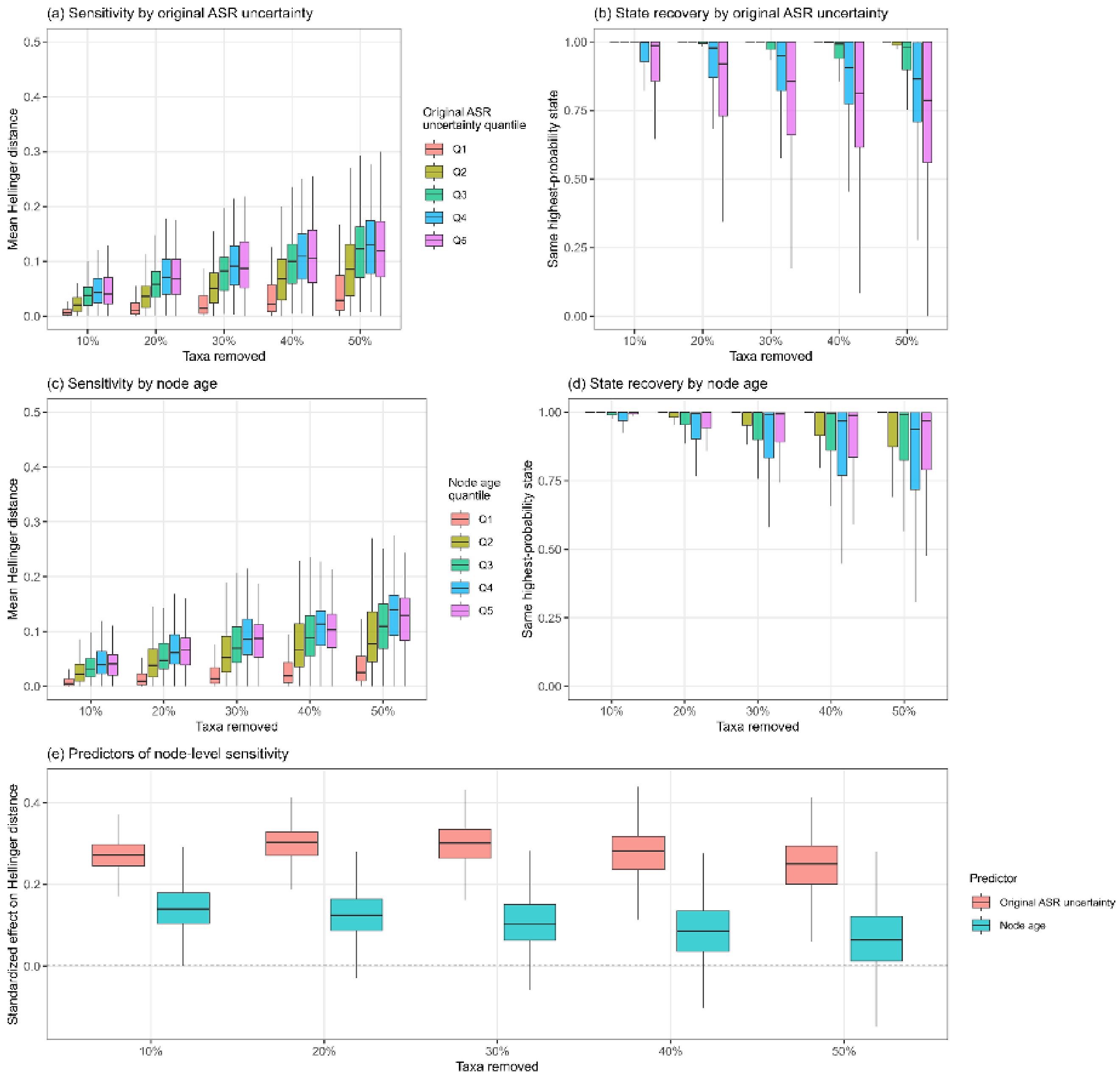

(a) Sensitivity by original ASR uncertainty
Mean Hellinger distance
Taxa removed
10%
20%
30%
40%
50%
Original ASR uncertainty quantile
Q1
Q2
Q3
Q4
Q5
(b) State recovery by original ASR uncertainty
Same highest-probability state
Taxa removed
(c) Sensitivity by node age
Mean Hellinger distance
Taxa removed
Node age quantile
Q1
Q2
Q3
Q4
Q5
(d) State recovery by node age
Same highest-probability state
Taxa removed
(e) Predictors of node-level sensitivity
Standardized effect on Hellinger distance
Taxa removed
Predictor
Original ASR uncertainty
Node age

**Figure S6: Correlated evolution between biome or dispersal syndrome and fleshiness or dehiscence in angiosperms.**

Transition rates based on the best fitting models for correlated evolution are shown, with arrow widths scaled to transition rates. (a) rates of biome shifts are dependent on dehiscence. The arrows in the middle represent transitions between dispersal syndromes regardless of biome. (b) evolution of fleshiness and biome are interdependently correlated. (c) fleshiness and zoochorous dispersal are interdependently correlated. (d) dehiscence and zoochorous dispersal are interdependently correlated. (e) fleshiness and dehiscence are interdependently correlated. Rates are in transitions Myr$^{-1}$. Dashed arrows indicate a transition rate <0.0005 transitions Myr$^{-1}$. *Not to scale as rates were too high for visualisation.

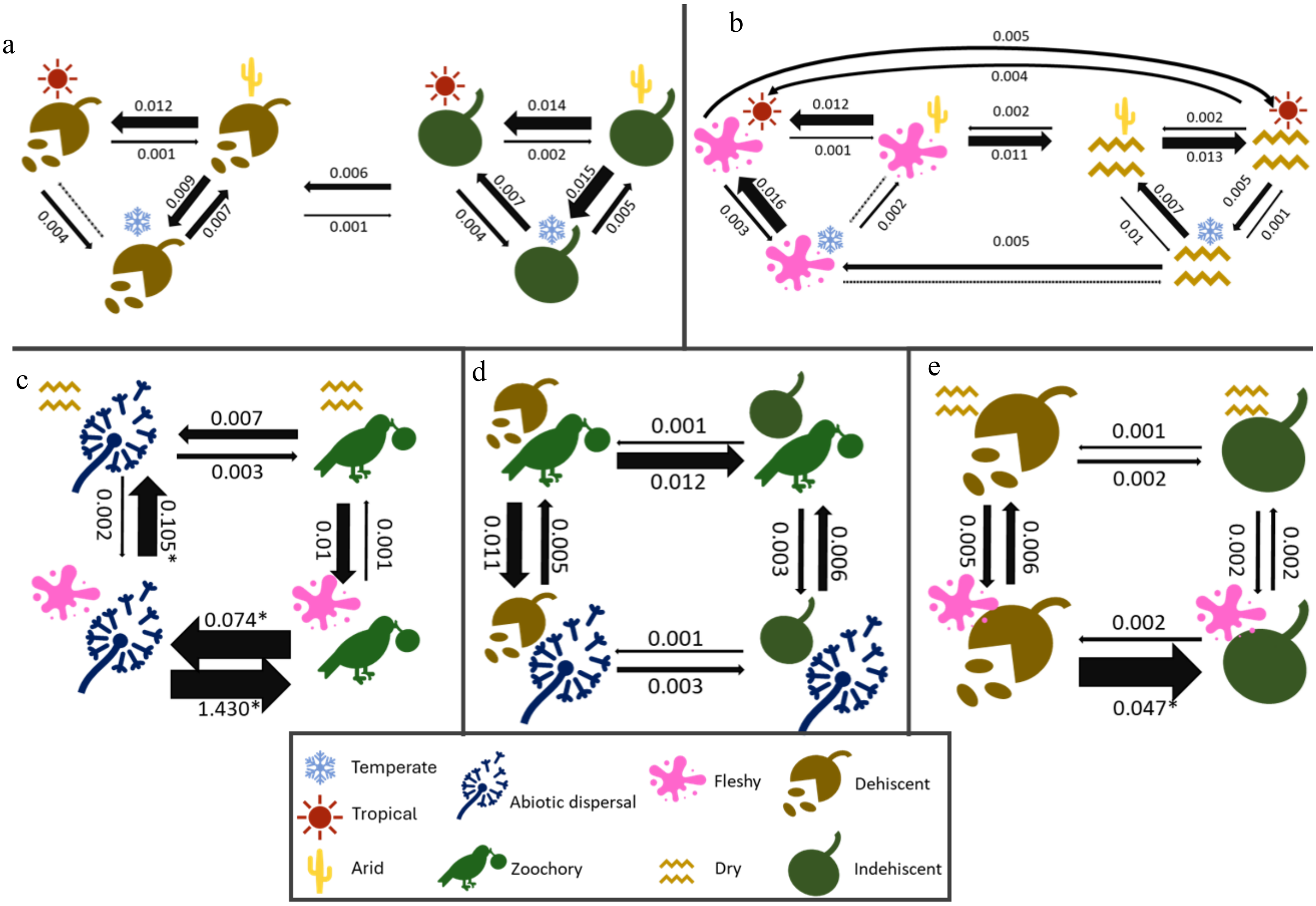

Table S1: Definitions of characters collected and the corresponding character states.

| Character | Definition and classification |
| --- | --- |
| **Diaspore type** | Functional dispersal unit for seed dispersal. Categorized as: whole fruit, seed, covered seed (seed at least partially covered with some sort of functional structure, e.g., an aril or an elaiosome), or floral fruit (diaspore includes a persistent functional floral organ, such as a fleshy calyx, a wing-like bract, or a fleshy receptacle). |
| **Dehiscence** | Whether or not the true fruit is truly dehiscent, so excluding schizocarp fruits. |
| **Fleshiness** | Whether or not the true fruit has a fleshy pericarp. |
| **Nutrition** | Absence or presence of any nutritious structure e.g. elaiosome, fleshy pulp, fleshy aril, fatty seed with diaspore. |
| **Released** | Whether or not the diaspore is intentionally released from the plant (so fruits that drop when they start to rot do not count, but fruits or seeds released consistently at or slightly before ripening do). |
| **Fruit size** | Average reported fruit size in centimetres (including possible functional components from the calyx or corolla) (along longest axis) |
| **Seed size** | Average reported seed size in millimetres (along longest axis) |
| **Proportional seed size** | = seed size/ (10*fruit size) In order to roughly approximate the number of seeds, as fruits with proportionally smaller seeds tend to have a greater number and vice versa. |

Table S2: Dispersal syndrome classification as used in this study.

Dispersal syndromes, the most relevant subtypes, and the defining diaspore characters were scored for 1201 species. We followed the most prevalent dispersal syndromes as defined in Beckman & Sullivan (2023).

| **Dispersal syndrome** | **Subtype** | **Definition** | **Key dispersal-related traits and diaspore characters** |
|---|---|---|---|
| Zoochory | Endozoochory | Diaspores eaten and dispersed by animals after passing digestive tract | Diaspore with fleshy nutritious structure, often conspicuous and/or pungent |
| | Synzoochory | Diaspores collected by animals and subsequently lost or forgotten about by disperser | Diaspore often large with nutritious seed, released, direct observation preferred |
| | Epizoochory | Diaspores dispersed by temporarily attaching to an animal | Diaspore sticky, or with hook- or needle-like appendages |
| | Myrmecochory | Diaspores dispersed by ants | Diaspore with elaiosome, often with targeted chemical cues |
| Anemochory | Flyers | Diaspores dispersed by gliding with drafts of wind | Winged, dust-like, or feathery diaspore |
| | Chamaechory | Diaspores or whole plant organs are released and blown by wind before seed release | Dry, lightweight, detaching parent plant(-part), released |
| Hydrochory | Nautochory | Diaspores that float on bodies of water | Corky or partially hollow, buoyant, diaspore |
| Autochory *sensu stricto* | Ballistic | Diaspores ejected by elastic energy stored in fruit or floral tissue | Explosively dehiscent fruit, lightweight seeds |
| Barochory (autochory *sensu lato*) | Unassisted dispersal | Diaspores released passively without further adaptations for dispersal | Diaspore released, no obvious adaptations |
| Antitelochory (autochory *sensu lato*) | Synaptospermy | Diaspores prevented from dispersal by attachment to parent or substrate | Diaspore unreleased or mucilaginous |
| | (Hypo)geocarpy | Diaspores produced or deposited at or under soil level by parent | Diaspore produced at ground level or underground |

### Table S3: Loadings of the first three principal component axes.

Component scores of the first three principal components. Percentages indicate amount of variance explained by each principal component.

| Character | PC1 (38.74%) | PC2 (16.30%) | PC3 (13.18%) |
|---|---|---|---|
| **Fleshiness** | 0.3956122 | -0.13833038 | -0.23836686 |
| **Dehiscence** | -0.3605368 | -0.40192034 | -0.16339928 |
| **Nutritious structure** | 0.3971244 | -0.26159066 | -0.14199760 |
| **log(Fruit size)** | 0.1259927 | -0.49397686 | 0.37207943 |
| **log(Seed size)** | 0.2574387 | -0.26411802 | 0.44226641 |
| **Diaspore = seed** | -0.3985812 | -0.13739297 | -0.18605845 |
| **Diaspore = fruit** | 0.3700131 | 0.22480723 | -0.27920175 |
| **Diaspore = covered seed** | 0.03701786 | -0.47574338 | 0.03185759 |
| **Diaspore = floral fruit** | 0.00009760 | 0.28030697 | 0.63789221 |
| **Diaspore released** | -0.3810950 | 0.05190198 | 0.16419990 |
| **log(Seed proportion)** | 0.1768253 | 0.24517527 | 0.12174389 |

Table S4: Summary of the Mk-models fitted for each character and data partition. Model with the lowest AICc in bold.

| Character | Models tested | No. of parameters | lnLikelihood | AICc | DeltaAICc | Weight | No. optimization attempts | Ancestral state (marginal likelihood) |
|---|---|---|---|---|---|---|---|---|
| Dispersal syndrome (anemochory, autochory, hydrochory, zoochory) | ER | 1 | -1355.771 | 2713.545 | 186.784 | 0 | 30 | Zoochory (99.3%) |
| | SYM | 6 | -1267.877 | 2547.824 | 21.064 | 0 | 30 | Zoochory (98.5%) |
| | CUSTOM | 6 | -1277.125 | 2566.319 | 39.559 | 0 | 30 | Hydrochory (67.5%) |
| | **ARD** | **12** | **-1251.25** | **2526.76** | **0** | **0.9296** | **30** | **Zoochory (97.3%)** |
| | HiddenER | 3 | -1348.888 | 2703.795 | 177.035 | 0 | 60 | Zoochory (99.6%) |
| | HiddenSYM | 13 | -1257.27 | 2540.844 | 14.084 | 0.0008 | 60 | Zoochory (99.7%) |
| | HiddenARD | 26 | -1239.379 | 2531.947 | 5.186 | 0.0695 | 60 | Hydrochory (94.6%) |
| Fleshiness (dry, fleshy) | ER | 1 | -659.804 | 1321.612 | 26.581 | 0 | 20 | Fleshy (92.4%) |
| | ARD | 2 | -657.995 | 1320 | 24.969 | 0 | 20 | Fleshy (99.4%) |
| | HiddenER | 3 | -647.964 | 1301.948 | 6.917 | 0.0305 | 20 | Fleshy (81.7%) |
| | **HiddenARD** | **6** | **-641.481** | **1295.031** | **0** | **0.9695** | **40** | **Fleshy (59.0%)** |
| Dehiscence (dehiscent, indehiscent) | ER | 1 | -665.726 | 1333.456 | 48.202 | 0 | 20 | Indehiscent (99.7%) |
| | ARD | 2 | -654.254 | 1312.518 | 27.265 | 0 | 20 | Indehiscent (98.4%) |
| | HiddenER | 3 | -651.384 | 1308.788 | 23.534 | 0 | 20 | Indehiscent (99.0%) |
| | **HiddenARD** | **6** | **-636.592** | **1285.254** | **0** | **1** | **40** | **Indehiscent (88.6%)** |
| Nutritious structure (absent, present) | ER | 1 | -689.259 | 1380.522 | 24.284 | 0 | 20 | Present (91.7%) |
| | ARD | 2 | -687.426 | 1378.863 | 22.625 | 0 | 20 | Present (98.0%) |
| | HiddenER | 3 | -678.214 | 1362.449 | 6.21 | 0.0429 | 20 | Present (78.6%) |
| | **HiddenARD** | **6** | **-672.084** | **1356.238** | **0** | **0.9571** | **40** | **Present (54.3%)** |
| Diaspore type (covered seed, floral fruit, fruit, seed) | ER | 1 | -1228.46 | 2458.923 | 169.34 | 0 | 20 | Fruit (99.9%) |
| | SYM | 6 | -1170.308 | 2352.685 | 63.103 | 0 | 20 | Fruit (99.5%) |
| | ARD | 12 | -1145.954 | 2316.169 | 26.586 | 0 | 20 | Fruit (89.2%) |
| | HiddenER | 3 | -1223.154 | 2452.329 | 162.746 | 0 | 40 | Fruit (99.8%) |
| | HiddenSYM | 13 | -1145.79 | 2317.883 | 28.301 | 0 | 40 | Fruit (99.7%) |
| | **HiddenARD** | **26** | **-1118.198** | **2289.583** | **0** | **1** | **40** | **Fruit (99.5%)** |

### Table S5: Mean transition numbers between dispersal syndromes throughout angiosperm evolutionary history.

Transitions based on the simmap analyses of dispersal syndrome ASR calculated over 1000 stochastic character maps. 95% high probability density interval of number of transitions is shown in brackets.

| **Transitions from ↓ / to →** | **anemochory** | **autochory** | **hydrochory** | **zoochory** |
|---|---|---|---|---|
| **anemochory** | | 62.80<br>(45–79) | 3.79<br>(1–7) | 128.52<br>(106–154) |
| **autochory** | 232.87<br>(201–260) | | 14.65<br>(9–21) | 176.35<br>(151–202) |
| **hydrochory** | 0.00<br>(0–0) | 34.63<br>(20–52) | | 0.00<br>(0–0) |
| **zoochory** | 0.00<br>(0–0) | 101.41<br>(84–122) | 30.59<br>(22–38) | |

Table S6: Summary of Mk-models fitted for each character combination tested for correlated evolution.

Model with the lowest AICc in bold for each character combination.

| Character comparison | Model tested | No. of parameters | lnLikelihood | AICc | DeltaAICc | Weight | No. iterations |
|---|---|---|---|---|---|---|---|
| Fleshiness / Dehiscence | independent | 4 | -1279.2407 | 2566.515 | 235.965628 | 0 | 20 |
| | **interdependent** | **8** | **-1157.2142** | **2330.549** | **0** | **0.8889** | **20** |
| | fleshiness is dependent | 6 | -1195.5239 | 2403.118 | 72.568984 | 0 | 20 |
| | dehiscence is dependent | 6 | -1161.3194 | 2334.709 | 4.159995 | 0.1111 | 20 |
| Dispersal / Pollination | independent | 4 | -861.6181 | 1731.273 | 2.646708 | 0.1586 | 20 |
| | interdependent | 8 | -857.766 | 1731.665 | 3.038503 | 0.1304 | 20 |
| | dispersal is dependent | 6 | -859.9187 | 1731.915 | 3.288344 | 0.1151 | 20 |
| | **pollination is dependent** | **6** | **-858.2745** | **1728.626** | **0** | **0.5958** | **20** |
| Dispersal / Biome | independent | 8 | -1742.4344 | 3500.998 | 32.671579 | 0 | 20 |
| | interdependent | 18 | -1715.8742 | 3468.369 | 0.042253 | 0.4124 | 20 |
| | dispersal is dependent | 12 | -1722.952 | 3470.186 | 1.858783 | 0.1663 | 20 |
| | **biome is dependent** | **14** | **-1719.9735** | **3468.327** | **0** | **0.4212** | **20** |
| Fleshiness / Biome | independent | 8 | -1674.7397 | 3365.609 | 44.978799 | 0 | 20 |
| | **interdependent** | **18** | **-1642.0047** | **3320.63** | **0** | **0.5233** | **20** |
| | fleshiness is dependent | 12 | -1649.2924 | 3322.866 | 2.2363 | 0.171 | 20 |
| | biome is dependent | 14 | -1646.7097 | 3321.799 | 1.169127 | 0.2916 | 20 |
| | only arid is dependent | 12 | -1651.7903 | 3327.862 | 7.232276 | 0.0141 | 20 |
| Dehiscence / Biome | independent | 8 | -1673.193 | 3362.516 | 10.725 | 0.0028 | 20 |
| | interdependent | 18 | -1659.19 | 3355.001 | 3.210172 | 0.1212 | 20 |
| | dehiscence is dependent | 12 | -1668.7489 | 3361.779 | 9.988834 | 0.0041 | 20 |
| | **biome is dependent** | **14** | **-1661.7054** | **3351.791** | **0** | **0.6032** | **20** |
| | only arid is dependent | 12 | -1664.5633 | 3353.408 | 1.617706 | 0.2687 | 20 |
| Fleshiness / Dispersal | independent | 4 | -1354.1578 | 2716.349 | 589.765453 | 0 | 20 |
| | **interdependent** | **8** | **-1055.2314** | **2126.584** | **0** | **0.8935** | **20** |
| | fleshiness is dependent | 6 | -1071.274 | 2154.618 | 28.034676 | 0 | 20 |
| | dispersal is dependent | 6 | -1059.3839 | 2130.838 | 4.25452 | 0.1065 | 20 |
| Dehiscence / Dispersal | independent | 4 | -1349.2312 | 2706.496 | 152.099598 | 0 | 20 |
| | **interdependent** | **8** | **-1269.1377** | **2554.396** | **0** | **0.6556** | **20** |
| | dehiscence is dependent | 6 | -1271.8066 | 2555.684 | 1.287369 | 0.3444 | 20 |
| | dispersal is dependent | 6 | -1282.0806 | 2576.231 | 21.835193 | 0 | 20 |
